# Graph Theoretical Design of Biomimetic Aramid Nanofiber Nanocomposites as Insulation Coatings for Implantable Bioelectronics

**DOI:** 10.1101/2020.12.28.424604

**Authors:** Huanan Zhang, Drew Vecchio, Ahmet Emre, Samantha Rahmani, Chong Cheng, Jian Zhu, Asish C. Misra, Joerg Lahann, Nicholas A. Kotov

**Author notes:** Corresponding authors: Huanan Zhang; Nicholas A. Kotov.

## Abstract

An insulation material combining crack and delamination resistance, flexibility, strong adhesion, and biocompatibility is vital for implantable bioelectronic devices of all types. Creating a material with the combination of all these properties is a particularly distinct challenge for implantable electrodes. Here we describe a nanocomposite material addressing these technological challenges based on aramid nanofibers (ANFs) whose unique mechanical properties are complemented by the epoxy resins with strong adhesion to various surfaces. The nanoscale structure of the ANF/epoxy nanocomposite coating replicates the nanofibrous organization of human cartilage, which is known for its exceptional toughness and longevity. The structural analogy between percolating networks of cartilage and ANF was demonstrated using Graph Theory (GT) analysis. The match of multiple GT indexes indicated the near identical organization pattern of cartilage and ANF/epoxy nanocomposite. When compared with the standard insulating material for bioelectronics, *Parylene C,* the ANF/epoxy nanocomposite demonstrates excellent interfacial adhesion, biocompatibility, and low inflammatory response. This study opens the possibility for the development of insulation materials suitable for different types of electronics for neural engineering and other biomedical applications. Also important, GT analysis makes possible structural characterization of complex biological and biomimetic materials.

## Introduction

Implantable microelectronic devices are crucial for advancing the diagnostics and therapies of many medical conditions.^1^ Cochlear implants,^2^ deep brain stimulation electrodes,^3^ and neural recording devices^4,5^ exemplify some of the major achievements in this area. Enabled by the advances in microfabrication technology and development of ultra-small electronic devices^6–9^ and concomitant development of new polymeric and nanostructured materials,^6,10^ a significant amount of research has been devoted lately to the engineering of flexible electronic circuits and biomedical microelectromechanical systems (bioMEMS) using polymer and nanocomposites substrates.^11–14^ The strong motivation for these studies stems from the many advantages that compliant bioelectronics possess in terms of chronic functionality and long-term biocompatibility.^15,16^ Additionally, implantable and epidermal electrode arrays with high density of electrodes have the advantage of targeting specific tissue regions, interrogating threedimensional brain areas, and improving the overall signal-to-noise ratio of the acquired data.^9,17^

However, the electrical components of electrode assemblies must be isolated from biological fluids with a thin layer insulation coatings, which often fails far too quickly in implants and other bioelectronics devices. The insulating coatings continue to be a hefty challenge in the design and implementation of bioelectronics and there are multiple reasons for these difficulties.^10^ *<u>First</u>,* biological electrolytes not only create short circuits but they also cause erosion of the plastic and metallic components^18^ of the electronic circuits. The coating must withstand long-term exposure to an array of enzymes and reactive species, while retaining non-toxicity and biocompatibility. *<u>Second</u>,* the insulating layers must withstand a wide range of mechanical deformations imposed on the implant, which includes mechanical stresses during the implantation/application. Note that chronic operations of these devices both in soft and hard tissues also involve multiple types of deformation that can increase over time. Interfaces between metal and plastic parts are particularly vulnerable to delamination under stress, which has been documented in multiple studies.^19–22^ *<u>Third</u>*, the mechanical properties of the entire device must be similar to those of the tissue in which they are implanted or in contact with to avoid inflammatory responses. For example, brain implants with high stiffness cause slow and progressive inflammation that can be resolved with compliant electrodes.^9,23–26^ Mechanical deformations are inevitable in a moving body and achieving integration of high toughness, adhesion strength, and non-porous morphology is far from trivial. *<u>Fourth</u>*, in a multi-electrode array, each electrode must function independently. Insulation materials must isolate each electrode and prevent “cross-talk” of the neighboring electrodes.^27^ In addition to low conductivity, the insulating coatings must have specific impedance and dielectric tensor that determine the electrical properties at specific frequencies where the devices operate. This task becomes particularly difficult for the high-density electrode arrays that require ‘designer’ composites as insulators. *<u>Fifth</u>*, the insulation layers are the bioelectronic components that have the largest area of direct contact with the surrounding biological tissue. The latest developments in the field indicate that these coatings must provide additional functionalities such as reduction of protein adsorption,^28^ drug release,^29–31^ antibacterial,^32,33^ inflammation mitigation,^34,35^ or gene/nucleotide delivery.^32,36^ Moreover, the need for these additional functions creates a multifunctional design challenge where the coatings need to withstand other processing conditions related to the preparation and optimization of these additional functionalities. Some of these methods can be fairly simple to implement, but others result in pinholes, delamination, and solvent-based erosion of the underlying films. This problem can become particularly acute when the insulating layer must be thin enough to accommodate other physical constrains of the implant.

Materials used for insulation coatings include Teflon^©37^ epoxy resins,^38^ polyimides^39^ and parylenes, with the latter being the most common for all bioelectronics devices. Parylene films are made by polymerization of *p*-xylylene derivatives using chemical vapor deposition (CVD) under optimized conditions to produce insulating coatings that are uniform in thickness, chemically inert, and highly resistive.^40,41^ To make a pinhole-free film, the materials need to be deposited at a thickness of at least 100-200 nm with common thicknesses in the range of 5-15μm.^42^ Sequential deposition of multiple layers needed, for example, for chronic implants and bioMEMS, results in the eventual development of cracks between the coatings.^43,44^ Advances in the syntheses of *p*-xylylene monomers allowed inclusion of reactive groups to promote interfacial bonding and hydrophilic functional groups for subsequent surface functionalization,^41,45^ but at the expense of mechanical properties of the coating itself. Furthermore, parylenes, Teflon^©^, and polyimides are known for their poor adhesion to metals, a primary cause of device failure.^46, 49^ Several different approaches have been developed to improve the adhesion of parylenes including crosslinking,^41^ high temperature, pressure treatment,^50^ surface modification,^51–54^ and mechanical anchoring with mushroom-shaped surface features.^49^ However, these approaches reduce other desirable properties including crack-resistance, long-term chemical robustness, and blood clot promotion. High temperature and pressure conditions as well as complex surface patterning also have adverse effects on other components of the electronic assembly.

The challenges encountered in the development of the insulation layers for implants and bioMEMS are similar to those that emerged in the past for other materials used in biomedical,^55,56^ energy,^57,57^ optical,^58^ and load-bearing applications.^55,59^ At the core of these challenges is the fundamental difficulty of creating materials that combine multiple contrarian properties. Interestingly enough, nearly all of these challenges are being addressed using nanoscale components assembled in structural patterns mimicking different biological structures.^60,61^ Following this conceptual pathway, we extend the biomimetic materials design approach to the design of insulating layers for biomedical devices. We exploited aramid nanofiber (ANF) composites,^62–64^ a new family of coatings that combine high flexibility, crack resistance, and electrical insulation, which replicates the structural organization of cartilage. By complementing the mechanical properties of ANF-based composites derived from Kevlar macrofibers with the strong adhesion of epoxy resin, a versatile insulation material for a variety of bioelectronics applications has been engineered. It was found that the adhesion properties of ANF/epoxy nanocomposite are superior to Parylene C and affords effective insulation of the corrugated surfaces of electrode arrays. Furthermore, ANF/epoxy nanocomposites demonstrated better biocompatibility and faster cell proliferation than Parylene C. Also important, the preparation of ANF/epoxy nanocomposite does not require a CVD process, which simplifies device manufacturing.

## Results and Discussion

The material known for both strong adhesion to corrugated surfaces, resilience to cracking, mechanical flexibility, and toughness is tissue cartilage. These properties are associated with a specific nanoscale structure and the macromolecular composition of cartilage can be described as a percolating network of self-assembled collagen nanofibers (**Figure 1A**).^65^ The fibers produce the *skeleton* of this biomaterial^66^ while complementary biopolymers (proteoglycans and glycoproteins) fill the space between the nanofibers. Such architecture enables effective stress transfer through the material via the collagen network providing the toughness, flexibility, and vibrational damping characteristic of cartilage. At the same time, both the collagen nanofibers and soft amorphous biopolymers provide multiple attractive interactions with other materials, even providing strong adhesion to uneven bone surface. The nanofibrous surface of tissue is also known for promoting cell adhesion and proliferation,^67,68^ which is another unifying requirement for both cartilage and the insulating layers of implants.

**Figure 1.**
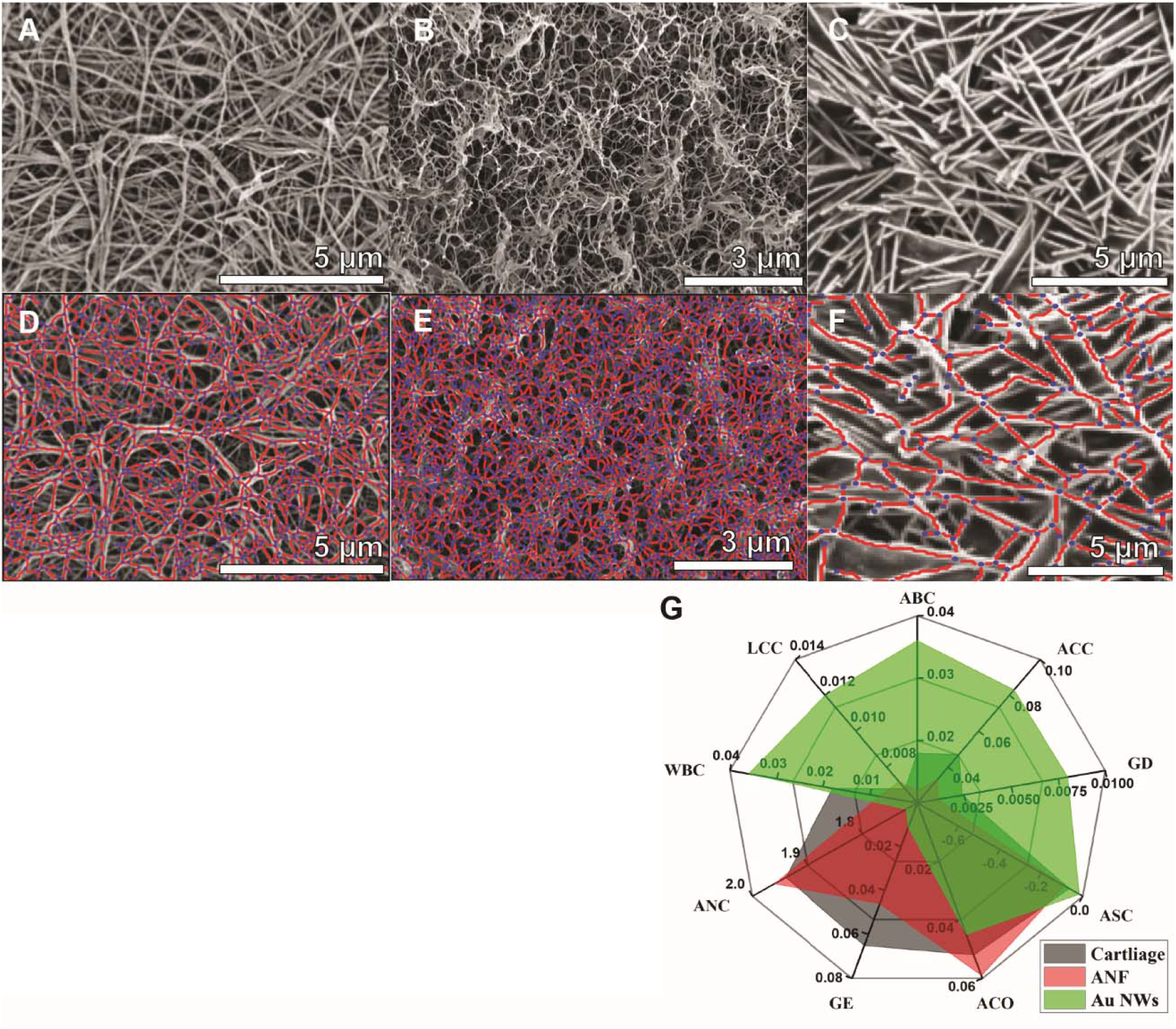
(A,D) SEM image of collagen nanofiber network from cartilage and its GT representation (D); SEM images of cartilage fibers was adapted from Ref.^66^ (B,E) SEM image of ANF network and its GT representation (E); (C,F) SEM image of gold nanowire network and its GT representation (F); SEM images of gold nanowires was adapted from Ref.^77^ The nodes in D-E are represented by red dots, and the edges are represented by green lines in the GT representations in **Figure 1**. Table 1 and spider plot (G) display the obtained GT descriptors for all three networks. Close overlap between cartilage and ANF networks can be observed.

Prior studies indicated that nanocomposites made from ANF nanofibers and common biocompatible polymers display mechanical properties with direct similarities to cartilage.^69^

Furthermore, visual assessment of the scanning electron microscopy images of collagen networks in cartilage^66^ and those of ANFs reveal distinct similarities (**Figure 1**). Unlike other biomimetic materials, for example nacre-like composites^70,71^ or silk-like polymer fibers,^72,73^ there is no established order parameter that can be directly associated with nanocomposite replicas of cartilage. To address this problem, we applied Graph Theory (GT) analysis of nanostructures^74,75^ to obtain direct quantitative evaluation of the similarity or dissimilarity of the nanofiber networks that form the skeletons of cartilage and ANF composites. Following the previous formalism,^75^ we describe the percolating network of nanofibers as graphs, *G*(*n,e*), where edges, *e*, represent nanofiber filaments, while nodes, *n*, represent their intersections. The network architecture can be described as a set of structural indices characterizing the structural patterns emerging in *G*(*n,e*). The GT analysis was carried out using software *StructuralGT*that converts SEM images into *G*(*n,e*).^75^ The indices calculated for percolating networks from collagen fibers and ANF obtained with similar magnification are given in **Table 1.** The detailed description of the indices used in the assessment of the materials are given in **Table 2.** Among them are the enumerators such as Average Degree (AD), Graph Density (GD), Average Clustering Coefficient (ACO), and Assortativity Coefficient (ASC)^76^ that assess short-range organization of the network represented, for instance, by branching at individual nodes. Long-range organization and connectivity of the nanofibrous structures is enumerated by Average Closeness Centrality (ACE) and Average Betweenness Centrality (ABC). Efficiency of the load transfer through the material is enumerated by Average Nodal Connectivity (ANC) and Global Efficiency (GE). Parameters such as Width-Weighted Average Betweenness Centrality (WBC) and Length-Weighted Average Closeness Centrality (LCC) provide scale-relevant enumeration of both structural and load transfer characteristics.

**Table 1.** Graph Theory Descriptor of Different Matetials

| Graph Theory Descriptor | Percolating Network Materials |  |  |
| --- | --- | --- | --- |
|  | Cartilage | ANF | Au NWs |
| Average Degree (AD) | 2.73 | 2.77 | 2.64 |
| Average Betweenness Centrality (ABC) | 0.018 | 0.012 | 0.036 |
| Average Closeness Centrality (ACC) | 0.047 | 0.033 | 0.083 |
| Graph Density (GD) | 0.0024 | 0.0011 | 0.008 |
| Assortativity Coefficient (ASC) | -0.065 | -0.09 | -0.016 |
| Average Clustering Coefficient (ACO) | 0.052 | 0.059 | 0.045 |
| Global Efficiency (GE) | 0.065 | 0.046 | 0.011 |
| Average Nodal Connectivity (ANC) | 1.94 | 1.96 | 1.72 |
| Width-weighted average Betweenness Centrality (WBC) | 0.018 | 0.008 | 0.036 |
| Length-weighted average closeness Centrality (LCC) | 0.0068 | 0.0072 | 0.012 |

**Table 2.**
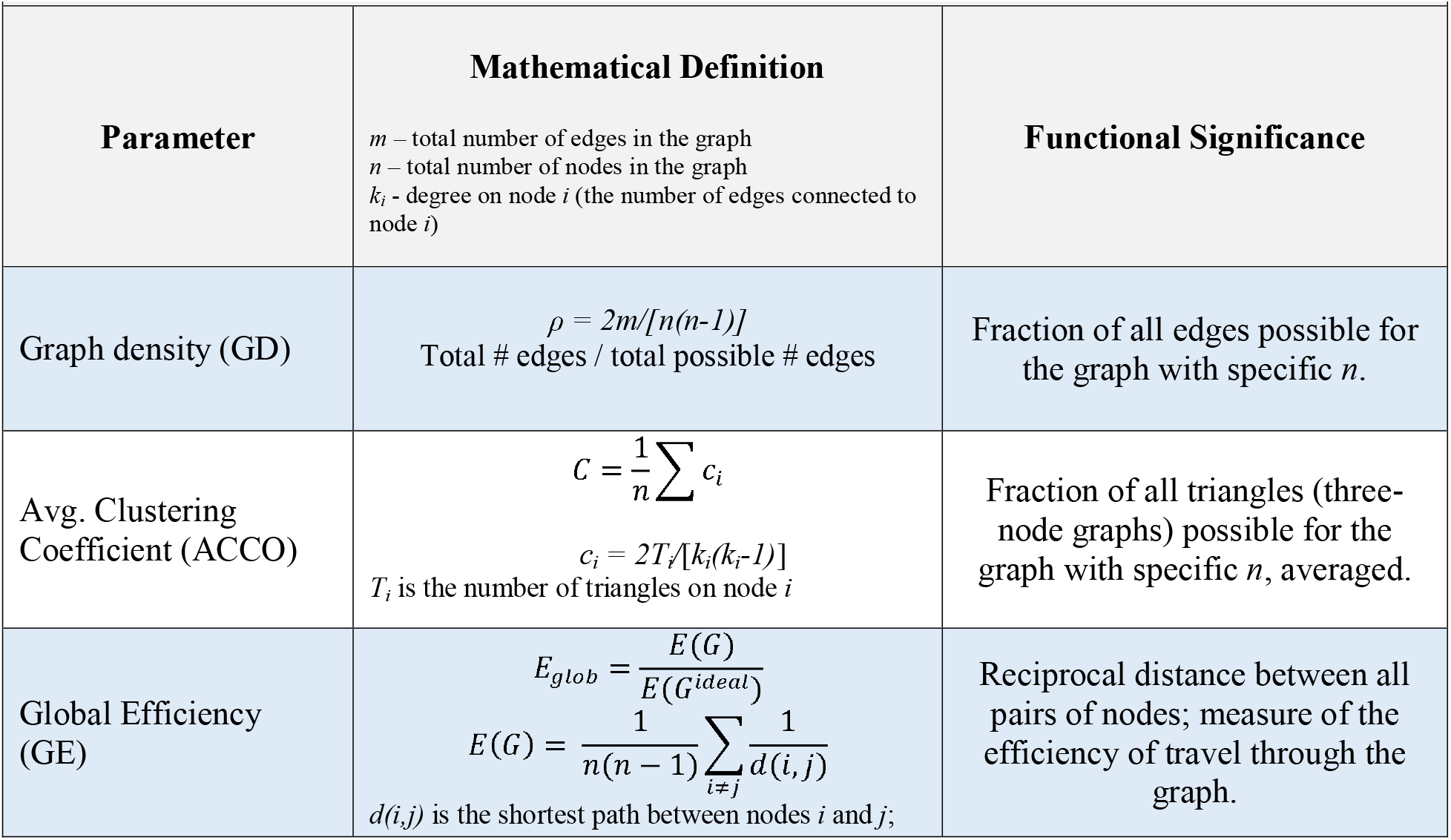
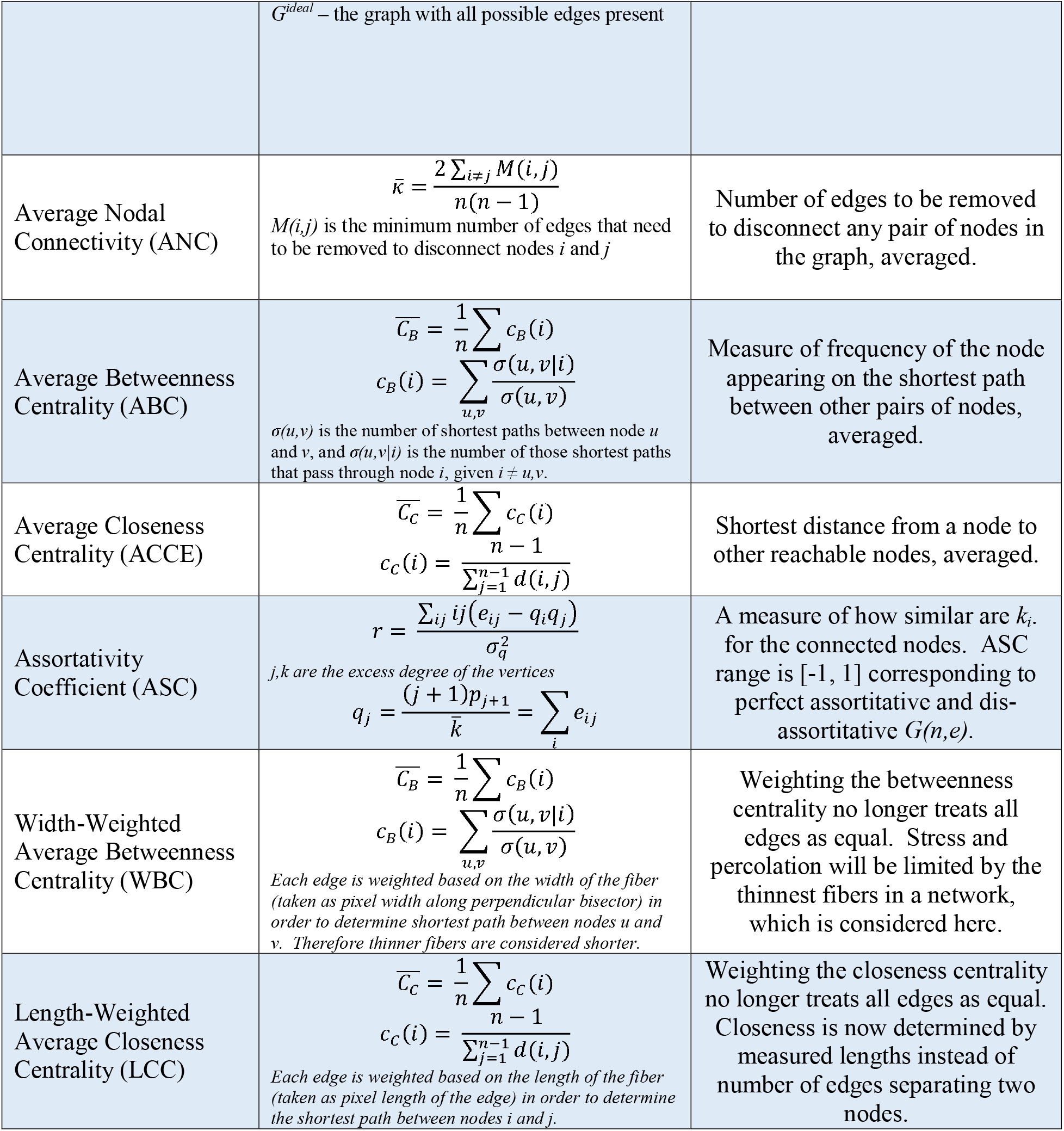
Graph theoretical enumerators applied for description of nanofiber networks and cartilage and ANF composites

Cartilage and ANF display a remarkable match across all the different GT parameters indicating pronounced similarity of their close- and long-range order, connectivity patterns, and load-transfer patterns. Note that this analysis does not include the physical characteristics of the load transfer, such the strength of the individual filaments, which will be different in cartilage and ANF. In this way, GT indices are analogous to the pattern descriptors being used for nacre and silk being most commonly described as materials based on a sequence of organic-inorganic layers and crystalline-amorphous polymer segments, respectively.

To verify the conclusions made with respect to cartilage and ANF, we applied the same analysis protocol to gold nanowire^77^ (NW) networks^77^ that serve in this study as a negative control for the index match for the ANF-cartilage pair. While having close values for AD, ACO and ASC, describing the short-range order, all other *G(n,e)* descriptors for Au NWs display distinct differences from both ANF and cartilage. Notably, the large differences in ACC and ANC, ACE, ABC, and GE highlight the vast differences in the in the long-range order and the load transfer between the networks made from NWs and those found for ANF and collagen.

The coatings from ANFs were produced by a spin coating process that allows control over the thickness of the deposited layers from 100 to 500 nm. Unlike Parylene C, no delamination between sequential ANF strata is observed. However, the produced ANF films are highly porous (**Figure 1B**), and in their native state would be very poor insulators. This apparent undesirable property can be turned into an advantage because high porosity and wettability of ANF layers allows other material components to be infiltrated into the network complementing ANF with additional functionalities. For example, the ANF network allows the infiltration of epoxy resin, which is known for superior adhesion, high strength, insulating properties, and biocompatibility. To briefly describe the process we used to prepare insulating layers (**Figure 2A)**, an ANF dispersion in DMSO was spin-coated onto a substrate, and then the DMSO was exchanged by water in a water bath. The excess water was then removed by an additional spin-coating cycle. An epoxy solution was then added in a similar fashion on top of the ANF film. The process is repeated in cycles to create films with desired thicknesses.

**Figure 2.**
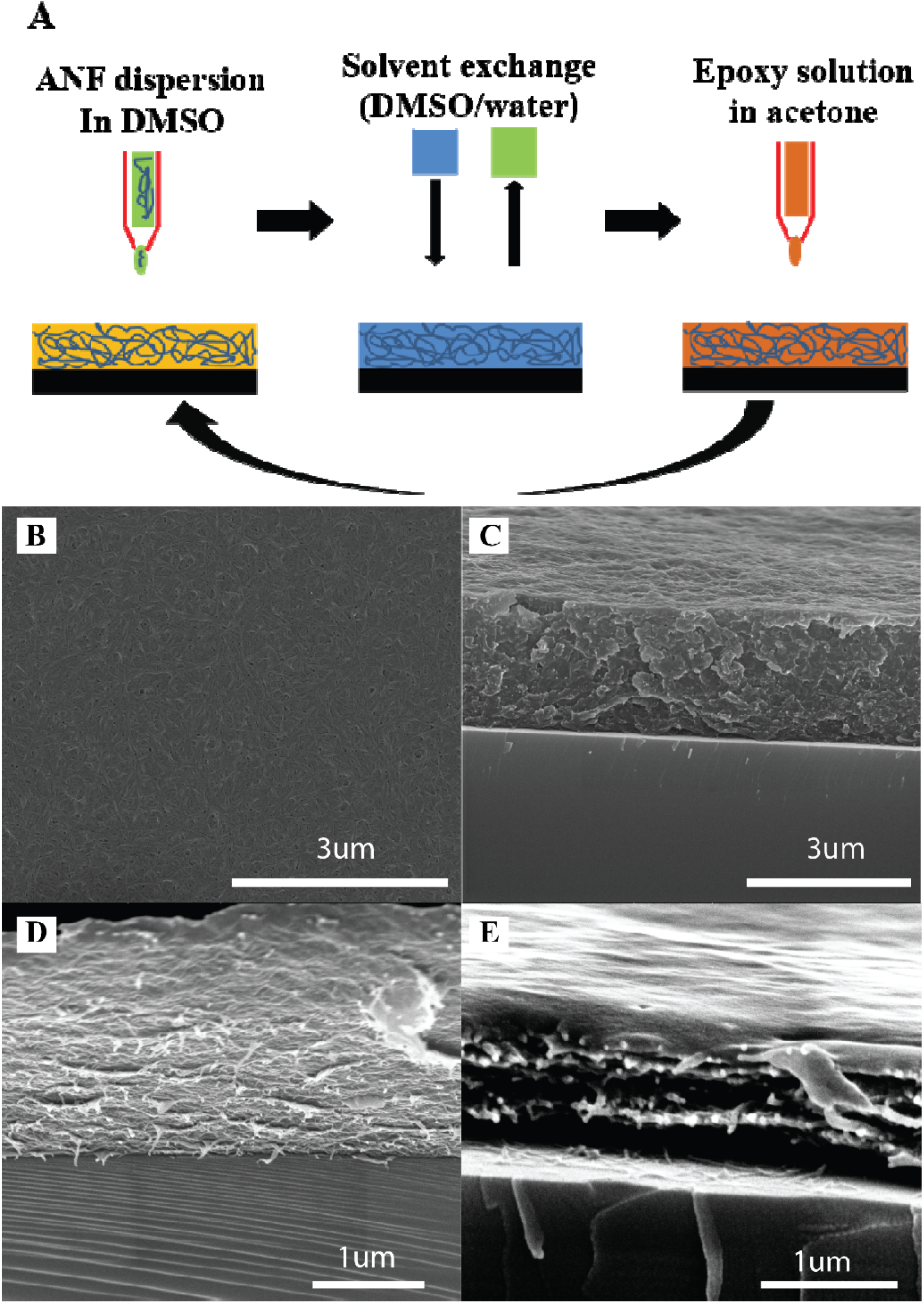
(A) Illustration of the nanocomposite coating process. (B) SEM image of ANF/epoxy composite after six deposition cycles. (C) SEM image of the cross-section of ANF/epoxy composite. (D) SEM image of the cross-section of layered ANF film without epoxy resin. (E) SEM image of the cross-section of the dried ANF film with epoxy resin.

From a cross-section of the scanning electron microscope (SEM) image of a six-bilayer ANF/epoxy nanocomposite, the average thickness is ~500 nm (**Figure 1B**). The surface of the ANF/epoxy coating was pinhole-free and uniform; it also displayed the nanoscale fibrous morphology expected from a thoroughly infiltrated nanocomposite (**Figure 2B**), which is crucial for insulating layers used in bioelectronics. The cross-sectional SEM image of the fractured coatings displayed fully filled layers without any pores or defects (**Figure 2C**).

Note that the films obtained from medical grade epoxy resin alone are deposited in the uniform films with layer thickness exceeding 100 μm due to poor wetting of the device surface.^78,79^ Furthermore, epoxy layers without supporting ANF network create defects at the protruding corners and edges of electrodes and other topographic features due to Maragoni effect^80,81^ resulting in increased surface roughness, coating thickness, and pinhole formation, which is particularly important for implantable electronic devices.^82^ Deposition of the nanofiber ‘skeleton’ alleviates both problems by soaking epoxy solution and conformally depositing fibrous network over the edges of electrodes.

Spin-coating the epoxy solution must be performed when water is still present in the ANF network, which at first, might seem somewhat counterintuitive. The water in the pores prevents the collapse of the network and allows the epoxy resin to penetrate between the nanofibers. To corroborate this point, the epoxy solution was spin-coated onto an ANF film after water was removed; this resulted in epoxy being on top of the ANF film, not within (**Figure 1E**). A SEM image of a cross-section of the six layers of ANF film without any epoxy solution (**Figure 2D**) showed that the thickness of the ANF film without infiltration was approximately five times smaller than that of the ANF/epoxy nanocomposite indicating again the collapse of the ANF network after water removal. This occurs because of strong capillary forces, which was confirmed by supercritical drying of the ANF film that preserves its structure very well (**Figure 1B**).

Once the ANF/epoxy nanocomposite was synthesized, our next objective was to compare the performance of the nanofibrous coatings to Parylene C, as packaging materials for implantable devices. In addition to bulk mechanical properties, two types of adhesive interactions contribute to this functionality: adhesion between electrode and the coating and cohesion between successive insulating layers. Importantly, both types of adhesive interactions are intrinsically coupled to bulk mechanical properties of the insulators because of the flexural deformations of the device during implantation and use.

To test the adhesion between electrode and insulator layers, both ANF/epoxy nanocomposite and Parylene C were deposited onto silicon substrates coated with triple metal layers of Cr/Au/Cr (20 nm/400 nm/20 nm). Photolithography was used to define an area on the insulators, and then ANF/epoxy or Parylene C layers were etched with O2 plasma to expose the top Cr layer underneath. After that, the test samples were incubated in phosphate-buffered saline (PBS) solution for 12 hours at room temperature, followed by immersion into Cr etchant for 30 seconds and subsequent observation under an optical microscope. If the adhesion between the insulator and metal was strong, the Cr etchant would only have limited access to the Cr surface and the apparent etch rate would be slow. If delamination of the insulation layer occurred, the Cr etchant would have greater access to the Cr surface and a fast etching rate would be observed. The optical microscopy images after Cr etching in **Figure 3** show that the etch rate for the Parylene C-coated substrate is much faster than for substrates protected by the ANF/epoxy composite.

**Figure 3.**
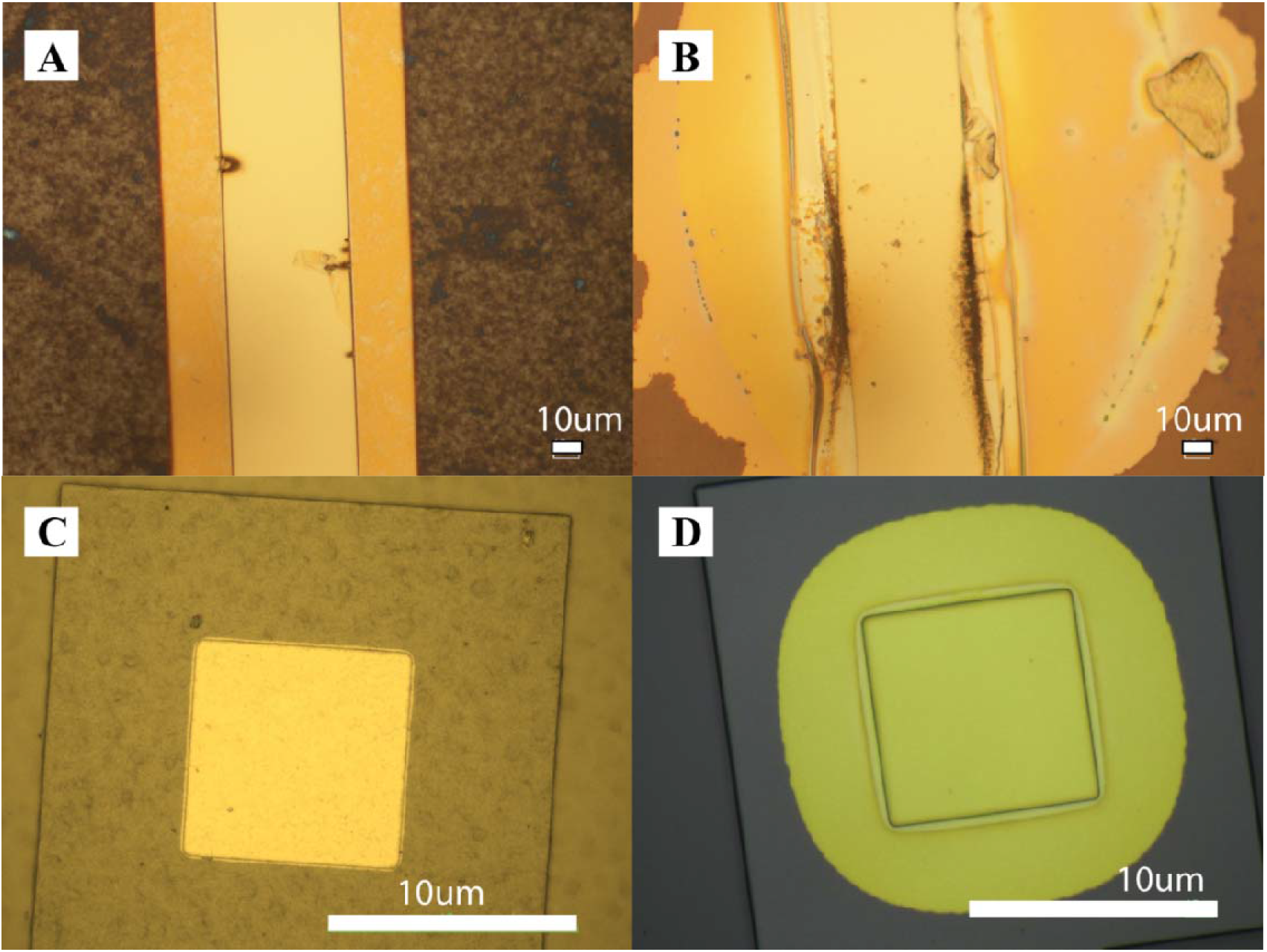
Metal adhesion test (A) Wet Cr etch of ANF/epoxy composite, rectangular strip opening. (B) Wet Cr etch of Parylene C, rectangular strip opening. (C) Wet Cr etch of ANF/epoxy composite, square opening. (D) Wet Cr etch of Parylene C, square opening.

Cohesion between successive insulating layers is essential because a typical implantable electrode device has metal microstructures sandwiched between two layers of insulators (**Figure 4A**). Thus, we prepared our interdigitated electrode (IDE) models in the same fashion as depicted in **Figure 4.** If the insulator-insulator interface has poor adhesion, crosstalk between the electrodes will increase and cause a dramatic decrease in the lateral impedance, Z_lat_. Because delamination of the metal-insulator interface can cause the Z_trans_ to decrease, the absolute value of Z_lat_ is a highly sensitive measurement for insulator/insulator interfacial adhesion (cohesion), especially for microstructures.

**Figure 4.**
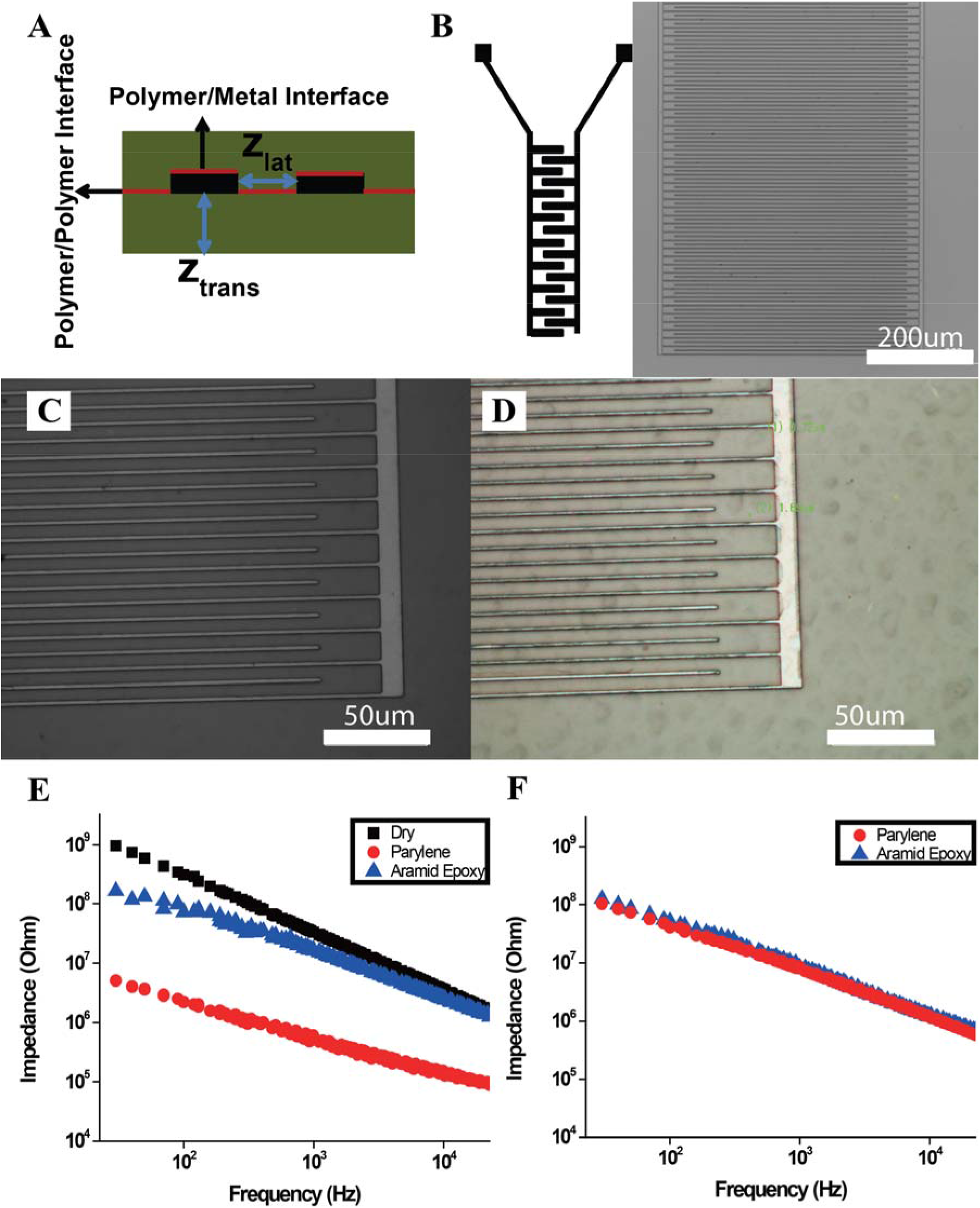
(A) Illustration of the interfaces in an interdigitated electrode (IDE). (B) Fabricated IDE structure. (C) Zoom in of IDE structure on Parylene C film. (D) Enlarged image of the IDE on ANF/epoxy composite. (E) Lateral impedance of different IDEs. (F) Transverse impedance of different IDEs.

To understand the compatibility of our ANF-epoxy nanocomposite with the photolithography processes used in manufacturing implantable electronics, conducting elements of the model IDE electrodes used in this study were similar to those used in high density electrode arrays (**Figure 4B**). Their lateral dimensions were 1.5 μm wide and 700 μm long with a high aspect ratio (**Figure 4B**, see Materials and Methods). Although ANF/epoxy has higher surface roughness compared to Parylene C, examination of the microelectrode structures by optical microscopy (**Figures 4C** and **4D)** shows that identical microscale structures analogous to high density electrode arrays can be fabricated on the ANF/epoxy nanocomposite as on the Parylene C substrates.

After an identical protocol to fabricate the IDE on both insulation materials was established, long-term soak tests were carried out to compare the insulator/insulator cohesion for Parylene C and the ANF/epoxy nanocomposite. After measuring the impedance of the electrodes in the dry state, 16 IDE electrodes of each insulator were soaked in PBS for 45 days in a temperature-controlled water bath at 37 °C (see Materials and Methods). Because there was no difference in dry Z_lat_ measurements for Parylene C and ANF/epoxy nanocomposite, only one dry Z_lat_ measurement was plotted in **Figure 4**. The impedance plot before the soaking time followed a linear relationship between impedance and frequency on a logarithmic scale (**Figure 4**), which indicated that the insulating layer performs as a simple resistor and capacitor circuit model. After the soak test, we observed the Z_lat_ of the Parylene C was one order of magnitude lower than that of the ANF/epoxy nanocomposite across the entire frequency spectrum. This finding demonstrated that IDEs insulated by Parylene C have more intense cross-talk than the IDEs insulated by our ANF/epoxy nanocomposite, which proves that our ANF/epoxy nanocomposite has stronger insulator/insulator cohesion than Parylene C does. In addition to Z_lat_, Z_trans_ was also measured to ensure the overall integrity of the IDE. Both IDE insulators displayed similar Z_trans_, which demonstrates that the difference in Z_lat_ arises from ongoing delamination at the insulator/insulator interface, not at the metal/polymer one, pointing to significant advantages of ANF/epoxy material for insulating coatings especially for complex electronic circuits and bioMEMS.

Finally, assessment of the cytotoxicity and biocompatibility of the new insulating coatings were carried out. NG108 neuroblastoma cells were cultured on the ANF/epoxy nanocomposite and the Parylene C substrates (**Figure 5**). The latter displayed limited cell attachment (**Figure 5B)** with the majority of the cells remaining suspended in the culture medium because Parylene C is too hydrophobic to support cell attachment and proliferation. On the contrary, NG108 cells can successfully attach and differentiate on the ANF/epoxy nanocomposite, demonstrating its biocompatibility, which was also vividly confirmed by Live/Dead assays (**Figures 5C** and **D**).

**Figure 5.**
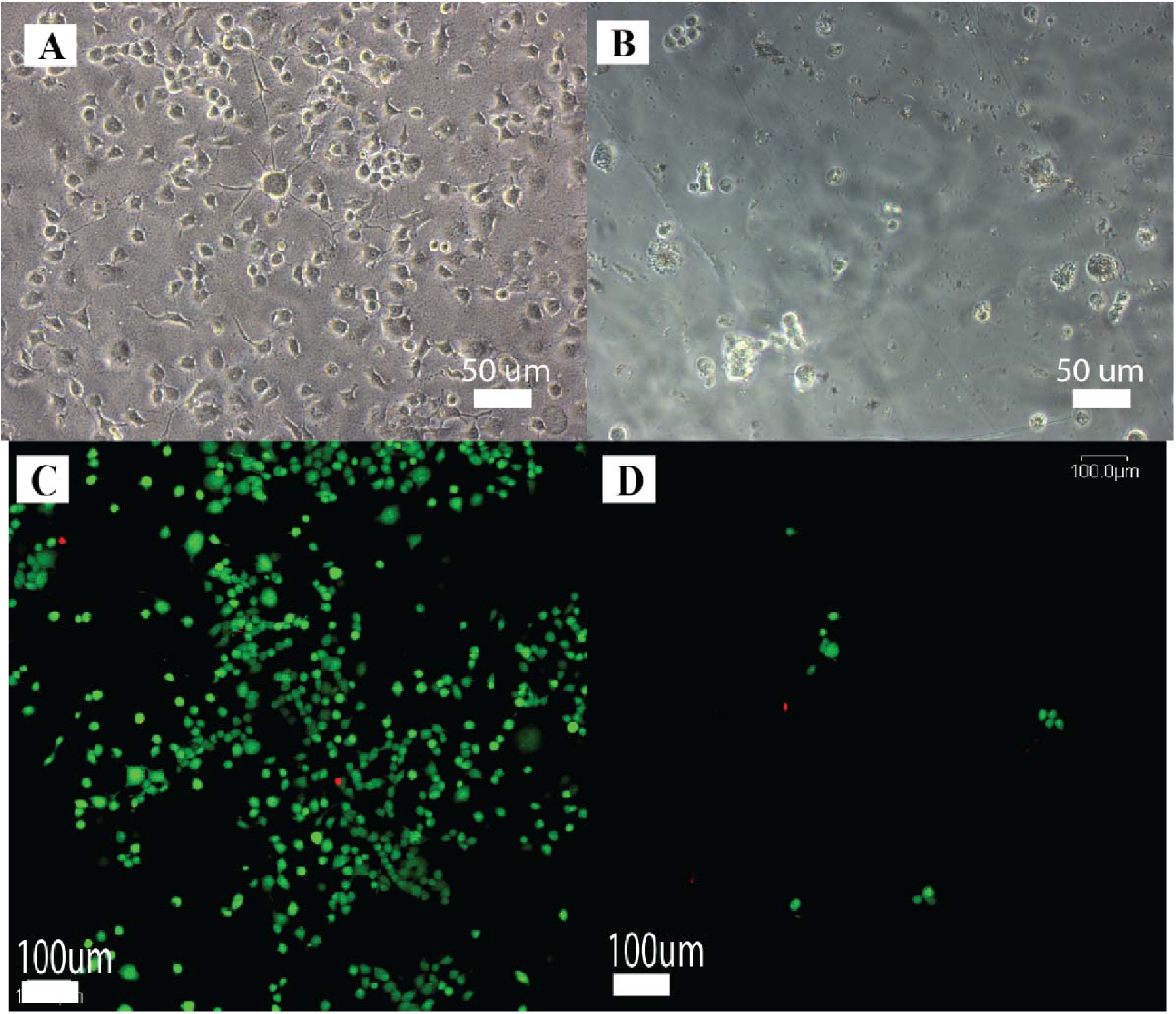
NG108 cell culture on various substrates. (A) NG108 cell culture on ANF/epoxy nanocomposite substrate. (B) NG108 cell culture on Parylene C. (C) Live/Dead assay of NG108 cell culture on ANF/epoxy (Green: Live, Red: Dead). (D) Live/Dead assay of NG108 cell culture on Parylene C (Green: Live, Red: Dead).

For chronic implants, platelet activation and thrombus formation could induce and amplify the inflammation response.^83^ Adhesion of platelets onto the insulation materials is the key event during thrombus formation that induces thrombosis, often leading to blood coagulation. Thus, the comprehensive characterization of the new nanocomposite coatings also required the assessment of platelet adhesion (**Figure 6)**. The platelets adhered to Parylene C exhibited irregular morphology and spreading pseudopodium. For the ANF/epoxy nanocomposite, the formation and spreading of pseudopodium was suppressed, resulting in a three-fold lower number of adhered platelets compared to Parylene C (**Figure 6C**).

**Figure 6.**
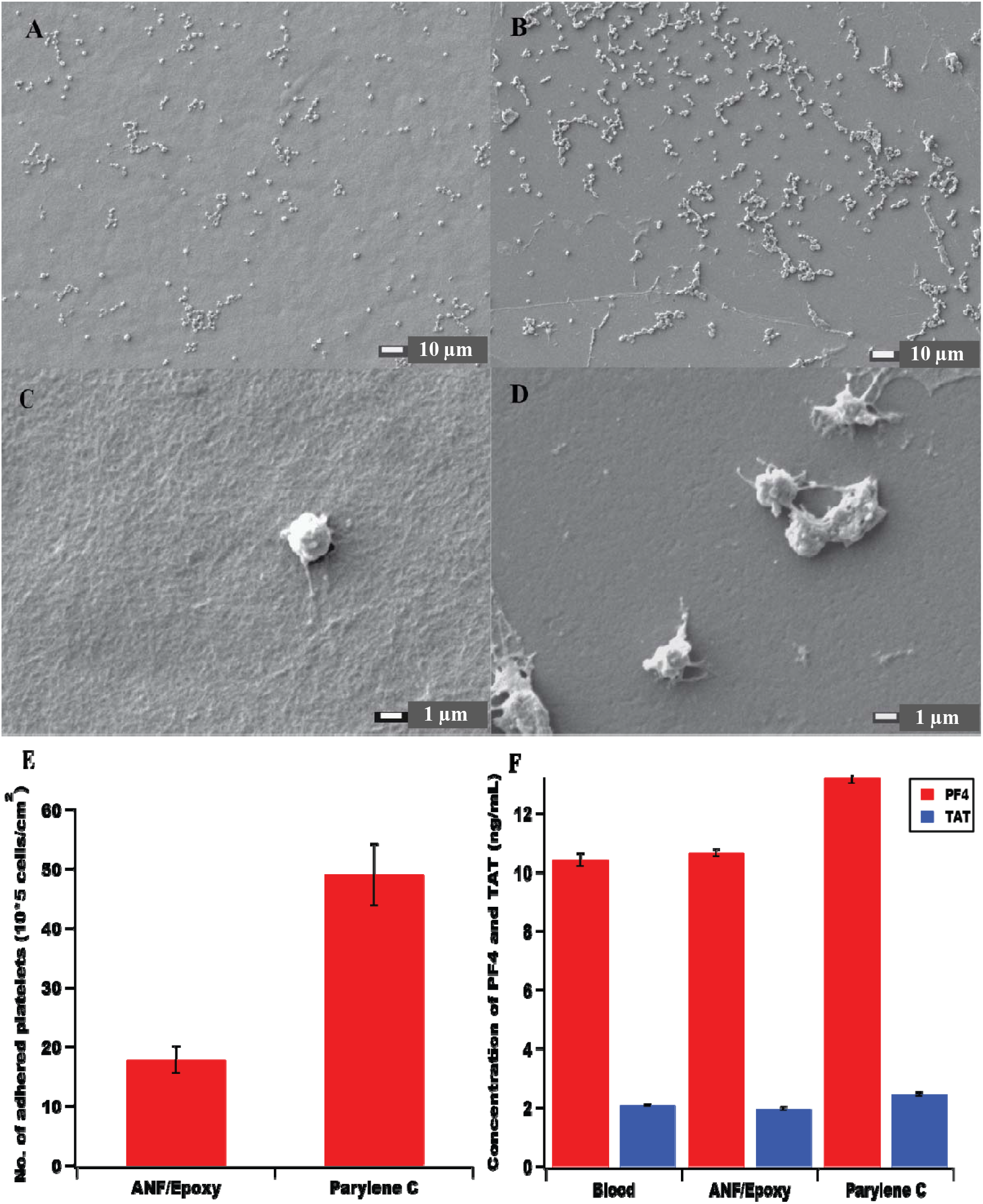
Blood compatibility of the insulation materials. (A) and (C). SEM morphology images of the platelets adhering to the ANF/epoxy nanocomposite. (B) and (D) SEM morphology images of the platelets adhering to the Parylene C film. (E) The average number of the adhered platelets onto the substrates from platelet-rich plasma estimated by 5 SEM images. (F) The generated Platelet Factor (PF-4) and thrombin–antithrombin (TAT) of the insulation substrates with 2 h blood incubation. The normal whole blood without exposure to insulation materials was applied as the control group. Values are expressed as means ± SD, n = 3.

The activation of platelets was further examined by measuring several signaling molecules. Platelet Factor 4 (PF4), also known as Chemokine Ligand 4, is released from alpha-granules of activated platelets during platelet aggregation. This process plays an important role in the promotion of blood coagulation and inflammation;^67^ therefore, PF4 was used to evaluate the platelet activation of our insulation materials. **Figure 6D** shows the PF4 concentrations in plasma after the two insulation materials being examined in this study were exposed to blood. The ANF/epoxy nanocomposite data show similar PF4 concentration as in normal whole blood, thus, no obvious platelet activation was observed. Parylene C showed an increased PF4 concentration and consequently, stronger platelet activation.

The contact activation of the coagulation system of the whole blood is another essential test for insulation materials when a certain amount of thrombin is generated during the activation of the coagulation cascade. The amount of thrombin was quantified via measurement of the thrombin-antithrombin III (TAT) level^84^ when the insulation materials came into contact with blood (**Figure 6D**). The concentration of TAT for the ANF/epoxy insulation coating was similar to that of whole blood, and considerably lower than that for Parylene C. Combining the results of the adhered platelets and the amounts of PF4 and TAT, one can conclude that the ANF/epoxy nanocomposite coating is more biocompatible than Parylene C, and is likely to lead to reduced inflammatory response.

## Conclusions

This study demonstrates that ANFs can be used as a platform to create biomimetic insulation materials opening up new opportunities for multifunctional material design. Enumeration of the network organization in the ANF/epoxy nanocomposite and collagen fibers in cartilage display a close match in multiple GT descriptors, such as AD, ABC, ACC, GD, ASC, ACO, GE, ANC, and LCC **(Table 1)**. From the fundamental perspective, we demonstrated here that GT analysis makes possible structural characterization of complex biological and biomimetic materials. Planar microelectronic structures can easily be created on ANF/epoxy nanocomposites and the results from the metal etching and long-term soak tests indicated that the ANF/epoxy nanocomposite displays better metal-insulator adhesion and insulator-insulator cohesion than Parylene C. In terms of biocompatibility, the neuroblastoma culture experiments validated the short-term biocompatibility of the new material. ANF/epoxy nanocomposite also exhibits lower blood activation, which could lead to a reduction of the inflammatory response. While having an auspicious set of properties, the ANF/epoxy nanocomposite needs to be tested in a long-term study, which is the next step in the development of this novel material.

## Materials and Methods

Right-hand twisted bulk Kevlar 69 was purchased from Thread Exchange. System 2000/2020 Epoxy Resin was purchased from FibreGlast Developments Corporation. All other chemicals were obtained from Sigma-Aldrich.

### ANF Nanofiber Dispersion

ANF dispersion was prepared using methods described elsewhere.^62^ Briefly, 1 g of aramid thread was dissolved in 100 mL of DMSO with 4 g of KOH to prepare a 10 mg/mL ANF nanofiber dispersion. The solution was stirred in a sealed container at room temperature for a week to ensure the complete dissolution of the nanofiber.

Supercritical CO_2_ drying (Leica CPD 300) was applied to ANF samples to prepare them for SEM (Thermo Fisher Helios 650 Nanolab SEM/FIB) imaging of the 3D network.

### Epoxy Solution

System 2000/2020 epoxy/hardener resin was mixed with a weight ratio of 3:1 as recommended by the manufacturer. After mixing, the epoxy resin mixture was added to acetone to make a 3% epoxy/acetone solution.

### ANF/Epoxy Nanocomposite

The nanocomposite was created through a spin-coating assisted deposition process. A 1 mL aliquot of as-prepared ANF nanofiber dispersion was spin-coated onto a 3.8 cm by 3.8 cm silicon oxide wafer at 1000 rpm. After spin coating, the film was rinsed with water to remove the excess DMSO. After rinsing with water, 1 mL of 3% epoxy resin in acetone was spin-coated on top of the ANF nanofiber film and dried in a standard oven at 100 °C for 1 min. The second layer of the epoxy resin solution was spin-coated onto the first layer and the layer-by-layer deposition process was repeated until the desired film thickness was achieved. Typically, six cycles result in a 3 μm thick film.

### Graph Theoretical Analysis

The GT analyses of the SEM microscopy images of ANF, collagen, and gold nanowire networks were performed by *StructuralGT,* a *Python* software package described in a previous publication.^75^ In brief, images were smoothed with a Gaussian blur to reduce salt-and-pepper noise, and the networks identified via OTSU thresholding. The resulting binary (B&W) images were skeletonized, serving as the basis for the graph representations of the original nanofiber networks *G(n,e).* Disconnected segments were removed from the skeleton, and the nodes and edges of the remaining graph were identified by the software. The final *G(n,e)* representations were plotted on top of the parent images using the package *MatPlotLib*^8^ The final *G(n,e)* representations obtained by *StructuralGT* were processed by the *NetworkX* library^86^ of algorithms to calculate the GT indices.

### Interdigitated Electrode (IDE) Fabrication

The procedure to fabricate interdigitated electrodes followed standard protocols in the field. Briefly, after assembling the bottom layer of the ANF/epoxy film, a positive photoresist (SPR220-3.0, Rohm, and Haas) was spin-coated to create the outline of the metal electrodes. Then Cr/Au/Cr (20 nm/400 nm/20 nm) was deposited by an E-beam evaporator (Enerjet Evaporator, Lesker) and lifted off in acetone. Then the top layer of the ANF/epoxy film was deposited on top of the liftoff metal structures. Positive photoresist (SPR-220-7.0, Rohm, and Haas) was spin-coated and developed again for the overall layout of the electrodes. Then the ANF/epoxy film was etched in oxygen plasma with 10% SF_6_ gas. Lastly, platinum microwire was attached on the bond pads of the electrodes using conductive silver epoxy. For Parylene C testing, 3 μm Parylene film was deposited in place of the ANF/epoxy film. Parylene C was etched in a pure oxygen plasma.

### Metal Adhesion Test

ANF/epoxy nanocomposite was first deposited on *e*-beam evaporated metals Cr/Au/Cr (20 nm/400 nm/20 nm). Then the ANF/epoxy film was etched in oxygen plasma with 10% SF6 gas to selectively open a square shape to expose the metals underneath. Parylene C film was etched in a pure oxygen plasma. To ensure experimental accuracy, both etched ANF/epoxy nanocomposite and Parylene C films were etched in Cr etchant for 30 s simultaneously, then rinsed in water for 5 min.

### Electrochemical Impedance Spectroscopy (EIS)

The impedance study was carried out on an Autolab PGSTAT 12; Frequency Response Analyzer software (EcoChemie, Utrecht, Netherlands) to record the impedance spectra of the electrodes. There were two electrodes in the interdigitated electrode design. For measurements of the interfacial impedance, the working electrode of the testing station was connected to one of the model implant electrodes. The counter/reference was connected to another electrode. An AC sinusoidal signal of 25 mV in amplitude was used to record the impedance over a frequency range of 10-32000 Hz. For transient impedance, the working electrode was connected to one of the electrodes. The counter electrode was connected to a gold foil immersed in phosphate-buffered saline (PBS), and a Ag/AgCl reference electrode was immersed in PBS.

### Long-Term Soaking Test

To measure long-term performance, both ANF/epoxy and Parylene C coated interdigitated electrodes were soaked in PBS at 37 °C. The impedance of the interdigitated electrodes was measured prior to the soaking test and measured again after soaking in PBS for 45 days.

### Scanning Electron Microscopy (SEM)

ANF/epoxy nanocomposite and Parylene C were deposited on a silicon wafer and coated with gold sputter for 60 s. SEM images were obtained using an FEI Nova Nanolab SEM at 10kV accelerating voltage.

### Blood Collection

Human blood sample was used to test the antithrombotic ability of the insulation materials. Fresh, whole human blood was collected by venipuncture from a healthy volunteer (male, 28 years old). The blood was mixed with 3.8 wt% anticoagulant citrate dextrose (9:1, v/v) and placed in a centrifuge at 1000 rpm and 4000 rpm for 15 min to obtain platelet-rich plasma (PRP). The PRP plasma and whole blood were stored at −20 °C until use.^87^

### Platelet Adhesion

The samples of ANF/epoxy nanocomposite- and Parylene C-coated glass slides (double sides coating, 1 × 1 cm^2^) were immersed in PBS (pH 7.4) and equilibrated at 37 °C for 1 h. After the removal of PBS, 1 mL of fresh PRP was introduced to each sample and incubated with PRP at 37 °C for 30 min. Then the PRP was decanted off, and samples were rinsed 3 times with PBS and finally treated with 2.5 wt% glutaraldehyde in PBS at 4 °C for 1 day. The prepared samples were then washed with PBS, and dehydrated by being passed through a series of graded alcohol/physiological saline solutions (50%, 70%, 80%, 90%, 95%, and 100 wt%, respectively; 15 min each) and isoamyl acetate/alcohol solutions (50%, 70%, 90%, 95%, and 100 wt%, respectively; 15 min each). The critical point drying of the substrates was done with liquid CO2. Platelet adhesion was investigated by SEM images. The numbers of adhered platelets on the insulation material surfaces were calculated from five SEM images.

### PF-4 and TAT Concentration Measurements

To evaluate the platelet activation and contact activation coagulation system, commercial enzyme-linked immunosorbent assays (ELISA) were used to test the generated factors in the blood. Platelet Factor 4 (PF-4) ELISA kits were purchased from Cusabio Biotech Co., Ltd, China; the thrombin-antithrombin III complex (TAT) ELISA kit, Enzygnost TAT micro, was purchased from Assay Pro, USA. For the test process, normal whole blood was incubated with the insulation materials for 2 h, and centrifuged for 15 min at 4 °C to obtain plasma. Then, the detections were carried out according to the respective instruction manuals as provided.

## Author contributions

H.Z. prepared ANF composites, performed characterization and co-wrote the paper; A.M., J.L. performed deposition of ParyleneC films and performed their characterization; A.E. prepared ANF composites for GT analysis; S.R. C.C., and J.Z assisted H. Z. in preparation and analyzed the performance of insulators; D.V. wrote the Python script for *StructuralGT* and carried out GT calculations of indexes; N.A.K developed GT analysis of biomimetic materials, conceptualized their analysis for composite characterization, and co-wrote the paper.

## References

1. Berger, T.W., Baudry, M., Brinton, R.D., Liaw, J.-., Marmarelis, V.Z., Park, A.Y., Sheu, B.J., and Tanguay, A.R. (2001). Brain-implantable biomimetic electronics as the next era in neural prosthetics. Proc. IEEE 89, 993–1012.

2. House, W.F. (1976). Cochlear implants. Ann Otol Rhinol Laryngol. 85, 1–93.

3. Perlmutter, J.S., and Mink, J.W. (2006). Deep brain stimulation. Annu. Rev. Neurosci. 29, 229–257.

4. Wise, K.D. (2005). Silicon microsystems for neuroscience and neural prostheses. IEEE Eng. Med. Biol. Mag. 24, 22–29.

5. Maynard, E.M., Nordhausen, C.T., and Normann, R.A. (1997). The Utah Intracortical Electrode Array: A recording structure for potential brain-computer interfaces. Electroencephalogr. Clin. Neurophysiol. 102, 228–239.

6. Kotov, N.A., Winter, J.O., Clements, I.P., Jan, E., Timko, B.P., Campidelli, S., Pathak, S., Mazzatenta, A., Lieber, C.M., Prato, M., et al. (2009). Nanomaterials for neural interfaces. Adv. Mater. 21.

7. Viventi, J., Kim, D.-H., Vigeland, L., Frechette, E.S., Blanco, J.A., Kim, Y.-S., Avrin, A.E., Tiruvadi, V.R., Hwang, S.-W., Vanleer, A.C., et al. (2011). Flexible, foldable, actively multiplexed, high-density electrode array for mapping brain activity in vivo. Nat. Neurosci. 14, 1599–1605.

8. Rodger, D.C., Fong, A.J., Li, W., Ameri, H., Ahuja, A.K., Gutierrez, C., Lavrov, I., Zhong, H., Menon, P.R., Meng, E., et al. (2008). Flexible parylene-based multielectrode array technology for high-density neural stimulation and recording. Sensors Actuators B Chem. 132, 449–460.

9. Yoshida Kozai, T.D., Langhals, N.B., Patel, P.R., Deng, X., Zhang, H., Smith, K.L., Lahann, J., Kotov, N.A., and Kipke, D.R. (2012). Ultrasmall implantable composite microelectrodes with bioactive surfaces for chronic neural interfaces. Nat. Mater. 11.

10. Sommakia, S., Lee, H.C., Gaire, J., and Otto, K.J. (2014). Materials approaches for modulating neural tissue responses to implanted microelectrodes through mechanical and biochemical means. Curr. Opin. Solid State Mater. Sci. 18, 319–328.

11. Hollenberg, B.A., Richards, C.D., Richards, R., Bahr, D.F., and Rector, D.M. (2006). A MEMS fabricated flexible electrode array for recording surface field potentials. J. Neurosci. Methods 153, 147–153.

12. Stieglitz, T., and Gross, M. (2002). Flexible BIOMEMS with electrode arrangements on front and back side as key component in neural prostheses and biohybrid systems. Sensors Actuators B Chem. 83, 8–14.

13. Kim, D.-H., Ahn, J.-H., Choi, W.M., Kim, H.-S., Kim, T.-H., Song, J., Huang, Y.Y., Liu, Z., Lu, C., and Rogers, J.A. (2008). Stretchable and Foldable Silicon Integrated Circuits. Science (80-.). 320, 507LP–511.

14. Kim, D.-H., Viventi, J., Amsden, J.J., Xiao, J., Vigeland, L., Kim, Y.-S., Blanco, J.A., Panilaitis, B., Frechette, E.S., Contreras, D., et al. (2010). Dissolvable films of silk fibroin for ultrathin conformal bio-integrated electronics. Nat. Mater. 9, 511–7.

15. Takeuchi, S., Suzuki, T., Mabuchi, K., and Fujita, H. (2003). 3D flexible multichannel probe array. In Proceedings of the IEEE Micro Electro Mechanical Systems (MEMS), pp. 367–370.

16. Polikov, V.S., Tresco, P.A., and Reichert, W.M. (2005). Response of brain tissue to chronically implanted neural electrodes. J. Neurosci. Methods 148, 1–18.

17. Wise, K.D., Angell, J.B., and Starr, A. (1970). An Integrated-Circuit Approach to Extracellular Microelectrodes. IEEE Trans. Biomed. Eng. BME-17, 238–247.

18. Bowman, L., and Meindl, J.D. (1986). The Packaging of Implantable Integrated Sensors. IEEE Trans. Biomed. Eng. BME-33, 248–255.

19. Bettinger, C.J. (2018). Recent advances in materials and flexible electronics for peripheral nerve interfaces. Bioelectron. Med. 4, 6.

20. Goding, J.A., Gilmour, A.D., Aregueta-Robles, U.A., Hasan, E.A., and Green, R.A. (2018). Living Bioelectronics: Strategies for Developing an Effective Long-Term Implant with Functional Neural Connections. Adv. Funct. Mater. 28, 1702969.

21. Ye, G., and Wang, X. (2010). Glucose sensing through diffraction grating of hydrogel bearing phenylboronic acid groups. Biosens. Bioelectron. 26, 772–7.

22. Simon, D.T., Gabrielsson, E.O., Tybrandt, K., and Berggren, M. (2016). Organic Bioelectronics: Bridging the Signaling Gap between Biology and Technology. Chem. Rev. 116, 13009–13041.

23. Zhang, H., Patel, P.R., Xie, Z., Swanson, S.D., Wang, X., and Kotov, N.A. (2013). Tissue-compliant neural implants from microfabricated carbon nanotube multilayer composite. ACS Nano 7, 7619–29.

24. He, F., Lycke, R., Ganji, M., Xie, C., and Luan, L. (2020). Ultraflexible Neural Electrodes for Long-Lasting Intracortical Recording. iScience 23, 101387.

25. Salatino, J.W., Ludwig, K.A., Kozai, T.D.Y., and Purcell, E.K. (2017). Glial responses to implanted electrodes in the brain. Nat. Biomed. Eng. 1, 862–877.

26. Patel, P.R., Zhang, H., Robbins, M.T., Nofar, J.B., Marshall, S.P., Kobylarek, M.J., Kozai, T.D.Y., Kotov, N.A., and Chestek, C.A. (2016). Chronic in vivo stability assessment of carbon fiber microelectrode arrays. J. Neural Eng. 13.

27. Wilke, R.G.H., Moghadam, G.K., Lovell, N.H., Suaning, G.J., and Dokos, S. (2011). Electric crosstalk impairs spatial resolution of multi-electrode arrays in retinal implants. J. Neural Eng. 8, 46016.

28. Onuki, Y., Bhardwaj, U., Papadimitrakopoulos, F., and Burgess, D.J. (2008). A review of the biocompatibility of implantable devices: current challenges to overcome foreign body response. J. Diabetes Sci. Technol. 2, 1003–15.

29. Takeshi, S., Greg, K., Shin-ichiro, H., R., B.L., Gerard, L., Robert, W., D., K.B., George, P., Pallassana, N., B., L.M., et al. (2001). Stent-Based Delivery of Sirolimus Reduces Neointimal Formation in a Porcine Coronary Model. Circulation 104, 1188–1193.

30. Scheller, B., Hehrlein, C., Bocksch, W., Rutsch, W., Haghi, D., Dietz, U., Böhm, M., and Speck, U. (2006). Treatment of Coronary In-Stent Restenosis with a Paclitaxel-Coated Balloon Catheter. N. Engl. J. Med. 355, 2113–2124.

31. Zilberman, M., and Elsner, J.J. (2008). Antibiotic-eluting medical devices for various applications. J. Control. Release 130, 202–215.

32. Deng, Z.J., Morton, S.W., Ben-Akiva, E., Dreaden, E.C., Shopsowitz, K.E., and Hammond, P.T. (2013). Layer-by-Layer Nanoparticles for Systemic Codelivery of an Anticancer Drug and siRNA for Potential Triple-Negative Breast Cancer Treatment. ACS Nano 7, 9571–9584.

33. Zhang, Z., Nong, J., and Zhong, Y. (2015). Antibacterial, anti-inflammatory and neuroprotective layer-by-layer coatings for neural implants. J. Neural Eng. 12 12, 046015.

34. He, W., McConnell, G.C., and Bellamkonda, R. V (2006). Nanoscale laminin coating modulates cortical scarring response around implanted silicon microelectrode arrays. J. Neural Eng. 3, 316–326.

35. He, W., McConnell, G., Schneider, T., and Bellamkonda, R. V (2007). A novel antiinflammatory surface for neural electrodes. Adv. Mater. 19, 3529–3533.

36. Kim, C.-H., Cha, S.-H., Kim, S.C., Song, M., Lee, J., Shin, W.S., Moon, S.-J., Bahng, J.H., Kotov, N.A., and Jin, S.-H. (2011). Silver nanowire embedded in P3HT:PCBM for high-efficiency hybrid photovoltaic device applications. ACS Nano 5.

37. Nicolelis, M.A.L., Dimitrov, D., Carmena, J.M., Crist, R., Lehew, G., Kralik, J.D., and Wise, S.P. (2003). Chronic, multisite, multielectrode recordings in macaque monkeys. Proc. Natl. Acad. Sci. 100, 11041 LP–11046.

38. Liu, X., McCreery, D.B., Carter, R.R., Bullara, L.A., Yuen, T.G.H., and Agnew, W.F. (1999). Stability of the interface between neural tissue and chronically implanted intracortical microelectrodes. IEEE Trans. Rehabil. Eng. 7, 315–326.

39. Stieglitz, T., Beutel, H., Schuettler, M., and Meyer, J.-U. (2000). Micromachined, Polyimide-Based Devices for Flexible Neural Interfaces. Biomed. Microdevices 2, 283–294.

40. Meng, E., and Tai, Y.-C. (2005). Parylene etching techniques for microfluidics and bioMEMS. In 18th IEEE International Conference on Micro Electro Mechanical Systems, 2005. MEMS 2005., pp. 568–571.

41. Seymour, J.P., Elkasabi, Y.M., Chen, H.-Y., Lahann, J., and Kipke, D.R. (2009). The insulation performance of reactive parylene films in implantable electronic devices. Biomaterials 30, 6158–6167.

42. Ortigoza-Diaz, J.., Scholten, K.., Larson, C.., Cobo, A.., Hudson, T.., Yoo, J.., Baldwin, A.., and Weltman Hirschberg, A.; Meng, E. (2018). Techniques and Considerations in the Microfabrication of Parylene C Microelectromechanical Systems. Micromachines 9, 422.

43. Chang, J.H., Lu, B., and Tai, Y. (2011). Adhesion-enhancing surface treatments for parylene deposition. In 2011 16th International Solid-State Sensors, Actuators and Microsystems Conference, pp. 390–393.

44. Ortigoza-Diaz, J., Scholten, K., and Meng, E. (2018). Characterization and Modification of Adhesion in Dry and Wet Environments in Thin-Film Parylene Systems. J. Microelectromechanical Syst. 27, 874–885.

45. Lahann, J. (2006). Vapor-based polymer coatings for potential biomedical applications. Polym. Int. 55, 1361–1370.

46. Yasuda, H., Yu, Q.S., and Chen, M. (2001). Interfacial factors in corrosion protection: an EIS study of model systems. Prog. Org. Coatings 41, 273–279.

47. Yamagishi, F.G. (1991). Investigations of plasma-polymerized films as primers for Parylene-C coatings on neural prosthesis materials. Thin Solid Films 202, 39–50.

48. Sharma, A.K., and Yasuda, H. (1982). Effect of glow discharge treatment of substrates on parylene substrate adhesion. J. Vac. Sci. Technol. 21, 994–998.

49. Liger, M., Rodger, D.C., and Tai, Y.-C. (2003). Robust parylene-to-silicon mechanical anchoring. In The Sixteenth Annual International Conference on Micro Electro Mechanical Systems, 2003. MEMS-03 Kyoto. IEEE, pp. 602–605.

50. von Metzen, R.P., and Stieglitz, T. (2013). The effects of annealing on mechanical, chemical, and physical properties and structural stability of Parylene C. Biomed. Microdevices 15, 727–735.

51. Zhang, Q., Phillips, H.R., Purchel, A., Hexum, J.K., and Reineke, T.M. (2018). Sustainable and Degradable Epoxy Resins from Trehalose, Cyclodextrin, and Soybean Oil Yield Tunable Mechanical Performance and Cell Adhesion. ACS Sustain. Chem. Eng. 6, 14967–14978.

52. Ramier, J., Boubaker, M. Ben, Guerrouache, M., Langlois, V., Grande, D., and Renard, E. (2014). Novel routes to epoxy functionalization of PHA-based electrospun scaffolds as ways to improve cell adhesion. J. Polym. Sci. Part A Polym. Chem. 52, 816–824.

53. Kim, P., Kim, D.H., Kim, B., Choi, S.K., Lee, S.H., Khademhosseini, A., Langer, R., and Suh, K.Y. (2005). Fabrication of nanostructures of polyethylene glycol for applications to protein adsorption and cell adhesion. Nanotechnology 16, 2420–2426.

54. Dolatshahi-Pirouz, A., Jensen, T., Kraft, D.C., Foss, M., Kingshott, P., Hansen, J.L., Larsen, A.N., Chevallier, J., and Besenbacher, F. (2010). Fibronectin Adsorption, Cell Adhesion, and Proliferation on Nanostructured Tantalum Surfaces. ACS Nano 4, 2874–2882.

55. Ferreira, A.D.B.L., Nóvoa, P.R.O., and Marques, A.T. (2016). Multifunctional Material Systems: A state-of-the-art review. Compos. Struct. 151, 3–35.

56. Tang, Z., Wang, Y., Podsiadlo, P., and Kotov, N.A. (2006). Biomedical Applications of Layer-by-Layer Assembly: From Biomimetics to Tissue Engineering. Adv. Mater. 18, 3203–3224.

57. Tatsidjodoung, P., Le Pierrès, N., and Luo, L. (2013). A review of potential materials for thermal energy storage in building applications. Renew. Sustain. Energy Rev. 18, 327–349.

58. Annadhasan, M., Basak, S., Chandrasekhar, N., and Chandrasekar, R. (2020). NextGeneration Organic Photonics: The Emergence of Flexible Crystal Optical Waveguides. Adv. Opt. Mater. 8, 2000959.

59. Lee, G.-H., Moon, H., Kim, H., Lee, G.H., Kwon, W., Yoo, S., Myung, D., Yun, S.H., Bao, Z., and Hahn, S.K. (2020). Multifunctional materials for implantable and wearable photonic healthcare devices. Nat. Rev. Mater. 5, 149–165.

60. Ha, N.S., and Lu, G. (2020). A review of recent research on bio-inspired structures and materials for energy absorption applications. Compos. Part B Eng. 181, 107496.

61. Zan, G., and Wu, Q. (2016). Biomimetic and Bioinspired Synthesis of Nanomaterials/Nanostructures.

62. Yang, M., Cao, K., Sui, L., Qi, Y., Zhu, J., Waas, A., Arruda, E.M., Kieffer, J., Thouless, M.D., and Kotov, N.A. (2011). Dispersions of aramid nanofibers: a new nanoscale building block. ACS Nano 5, 6945–6954.

63. Cao, K., Siepermann, C.P., Yang, M., Waas, A.M., Kotov, N.A., Thouless, M.D., and Arruda, E.M. (2013). Reactive aramid nanostructures as high-performance polymeric building blocks for advanced composites. Adv. Funct. Mater. 23.

64. Yang, M., Cao, K., Yeom, B., Thouless, M.D., Waas, A., Arruda, E.M., and Kotov, N.A. (2015). Aramid nanofiber-reinforced transparent nanocomposites. In Journal of Composite Materials, pp. 1873–1879.

65. Sharma, A., Licup, A.J., Jansen, K.A., Rens, R., Sheinman, M., Koenderink, G.H., and MacKintosh, F.C. (2016). Strain-controlled criticality governs the nonlinear mechanics of fibre networks. Nat. Phys. 12, 584–587.

66. Holder, A.J., Badiei, N., Hawkins, K., Wright, C., Williams, P.R., and Curtis, D.J. (2018). Control of collagen gel mechanical properties through manipulation of gelation conditions near the sol-gel transition. Soft Matter 14, 574–580.

67. Raimondo, T., Puckett, S., and Webster, T.J. (2010). Greater osteoblast and endothelial cell adhesion on nanostructured polyethylene and titanium. Int. J. Nanomedicine 5, 647–652.

68. Chen, N., Tian, L., Patil, A.C., Peng, S., Yang, I.H., Thakor, N. V, and Ramakrishna, S. (2017). Neural interfaces engineered via micro-and nanostructured coatings. Nano Today 14, 59–83.

69. Xu, L., Zhao, X., Xu, C., and Kotov, N.A.N.A. Water-Rich Biomimetic Composites with Abiotic Self-Organizing Nanofiber Network. Adv. Mater. 30, 1–6.

70. Tang, Z.Y., Kotov, N.A., Magonov, S., and Ozturk, B. (2003). Nanostructured artificial nacre. Nat. Mater. 2, 413–418.

71. Kotov, N.A., Dékány, I., and Fendler, J.H. (1996). Ultrathin graphite oxide-polyelectrolyte composites prepared by self □ assembly: Transition between conductive and non conductive states. Adv. Mater. 8, 637–641.

72. Mohammadi, P., Aranko, A.S., Landowski, C.P., Ikkala, O., Jaudzems, K., Wagermaier, W., and Linder, M.B. (2019). Biomimetic composites with enhanced toughening using silk-inspired triblock proteins and aligned nanocellulose reinforcements. Sci. Adv. 5, eaaw2541.

73. Zhang, W., Ye, C., Zheng, K., Zhong, J., Tang, Y., Fan, Y., Buehler, M.J., Ling, S., and Kaplan, D.L. (2018). Tensan Silk-Inspired Hierarchical Fibers for Smart Textile Applications. ACS Nano 12, 6968–6977.

74. Jiang, W., Qu, Z., Kumar, P., Vecchio, D., Wang, Y., Ma, Y., Bahng, J.H., Bernardino, K., Gomes, W.R., Colombari, F.M., et al. (2020). Emergence of Complexity in Hierarchically Organized Chiral Particles. Science (80-.). 368, 642–648.

75. Wang, M., Vecchio, D., Wang, C., Emre, A., Xiao, X., Jiang, Z., Bogdan, P., Huang, Y., and Kotov, N.A.N.A. (2020). Biomorphic structural batteries for robotics. Sci. Robot. 5, eaba1912.

76. Newman, M.E.J. (2003). Mixing patterns in networks. Phys. Rev. E 67, 26126.

77. Spain, E., McCooey, A., Joyce, K., Keyes, T.E., and Forster, R.J. (2015). Gold nanowires and nanotubes for high sensitivity detection of pathogen DNA. Sensors Actuators B Chem. 215, 159–165.

78. Voo, R., Mariatti, M., and Sim, L.C. (2011). Technique, Properties of epoxy nanocomposite thin films prepared by spin coating. J. Plast. Film Sheeting 27, 331–346.

79. Norrman, K., Ghanbari-Siahkali, A., and Larsen, N.B. (2005). 6 Studies of spin-coated polymer films. Annu. Reports Sect. “C” (Physical Chem. 101, 174–201.

80. Shiratori, S., and Kubokawa, T. (2015). Double-peaked edge-bead in drying film of solvent-resin mixtures. Phys. Fluids 27, 102105.

81. Mutyala, M.S.K., Javadi, A., Zhao, J., Lin, T.C., Tang, W., and Li, X. (2015). Scalable Platform for Batch Fabrication of Micro/Nano Devices on Engineering Substrates of Arbitrary Shapes and Sizes. Procedia Manuf. 1, 205–215.

82. Li, L., Zhang, G., Tuan, C., Moon, K., and Sun, R. (2016). Formation of Polymer Insulation Layer (Liner) on Through Silicon Vias (TSV) with High Aspect Ratio over 5:1 by Direct Spin Coating. In 2016 IEEE 66th Electronic Components and Technology Conference (ECTC), pp. 1713–1719.

83. Nie, S., Qin, H., Cheng, C., Zhao, W., Sun, S., Su, B., Zhao, C., and Gu, Z. (2014). Blood activation and compatibility on single-molecular-layer biointerfaces. J. Mater. Chem. B 2, 4911–4921.

84. Martinelli, I., De Stefano, V., and Mannucci, P.M. (2014). Inherited risk factors for venous thromboembolism. Nat. Rev. Cardiol. 11, 140–156.

85. Hunter, J.D. (2007). Matplotlib: A 2D graphics environment. Comput. Sci. Eng. 9, 99–104.

86. Hagberg, A., Schult, D., and Swart, P. (2008). Proceedings of the Python in Science Conference (SciPy): Exploring Network Structure, Dynamics, and Function using NetworkX. 11–15.

87. Cheng, C., Li, S., Nie, S., Zhao, W., Yang, H., Sun, S., and Zhao, C. (2012). General and Biomimetic Approach to Biopolymer-Functionalized Graphene Oxide Nanosheet through Adhesive Dopamine. Biomacromolecules 13, 4236–4246.

